## Supplementary Figure 1 for "Distinct cortical systems reinstate the content and context of episodic memories"

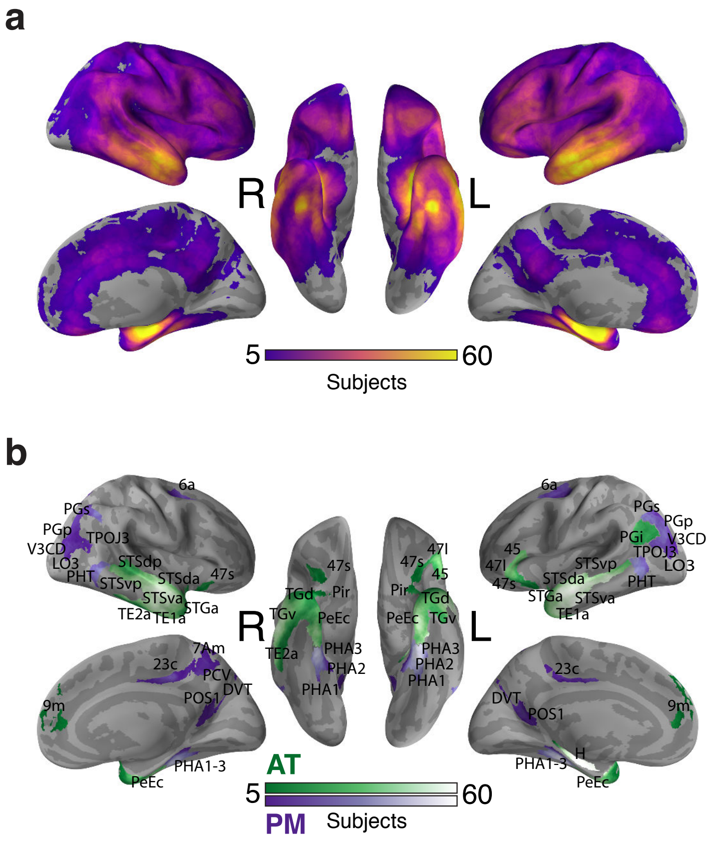


**Supplementary Figure 1   Electrode coverage.** **a**, Surface renders depict the number of subjects with electrode contacts within 10 mm of a vertex, after transformation into MNI space. **b**, The plot depicts the coverage as in **a**, highlighting parcels assigned to either the anterior temporal (AT) or posterior medial (PM) network. Parcel naming convention follows Glasser et al., 2016[30].
