## Supplementary Figure 2 for "Distinct cortical systems reinstate the content and context of episodic memories"

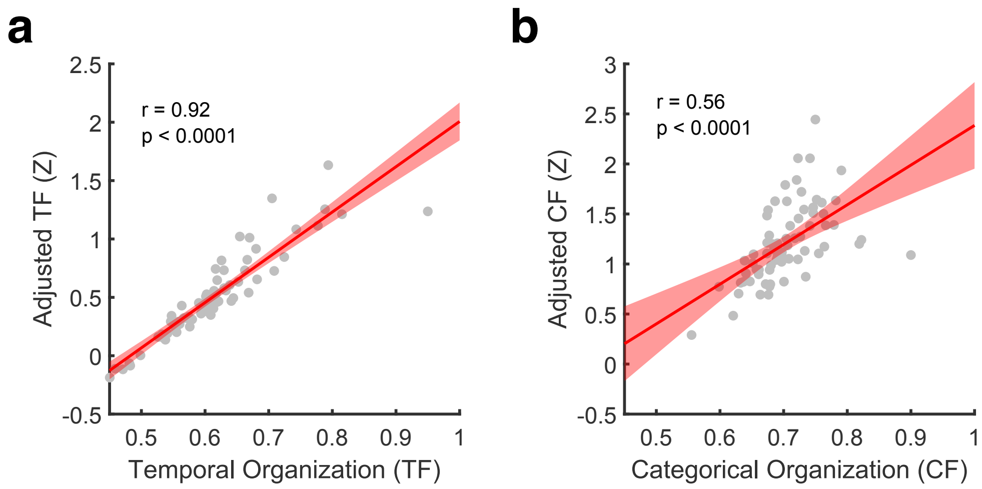


**Supplementary Figure 2 Measures of temporal and categorical organization are robust to list structure. a**, The scatter plot depicts the observed temporal factor (TF) score for each patient and the standardized TF score (in units of one standard deviation) after adjusting for the amount of temporal organization expected by chance from the category structure of each list. The red line indicates least squares linear fit across the group. **b**, The scatter depicts observed and adjusted category factor (CF) scores, accounting for the category structure of presented lists. Each point indicates a single subject. Shaded regions denote SEM ($n = 69$ subjects).
