## Supplementary Figure 3 for "Distinct cortical systems reinstate the content and context of episodic memories"

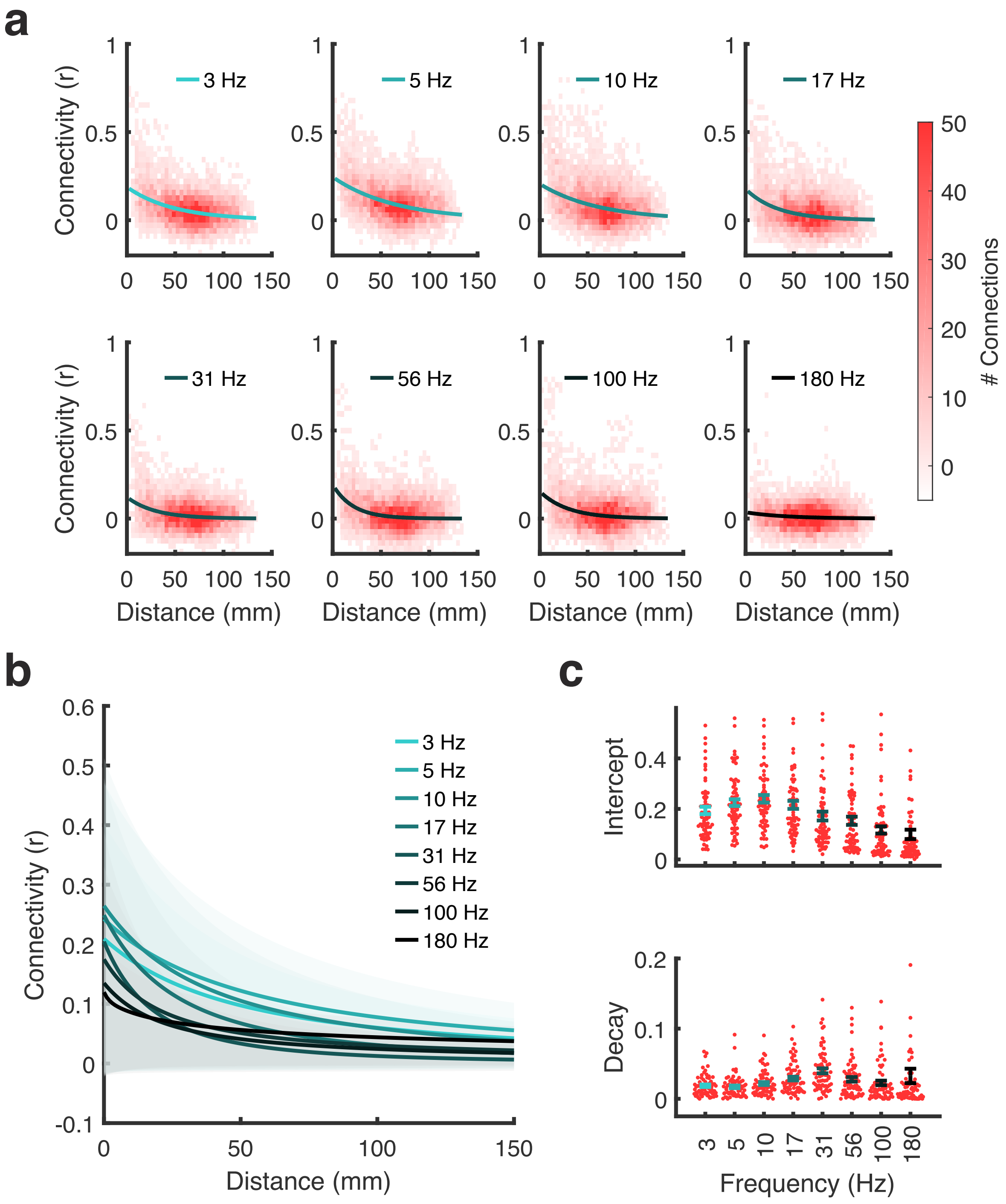


**Supplementary Figure 3 Connectivity between recording sites decreases with distance. a**, Plots depict intrinsic connectivity as a function of inter-electrode difference for an individual subject. Best fit exponentials are overlaid on each plot. **b**, Group-averaged ($n = 69$ subjects) predictions on the effect of inter-electrode distance on intrinsic connectivity. Shaded regions denote SEM. **c**, Lines depict group-averaged model parameters for describing the relation between distance and connectivity. Each point indicates a single subject. Error bars denote SEM.
