## Supplementary Figure 4 for "Distinct cortical systems reinstate the content and context of episodic memories"

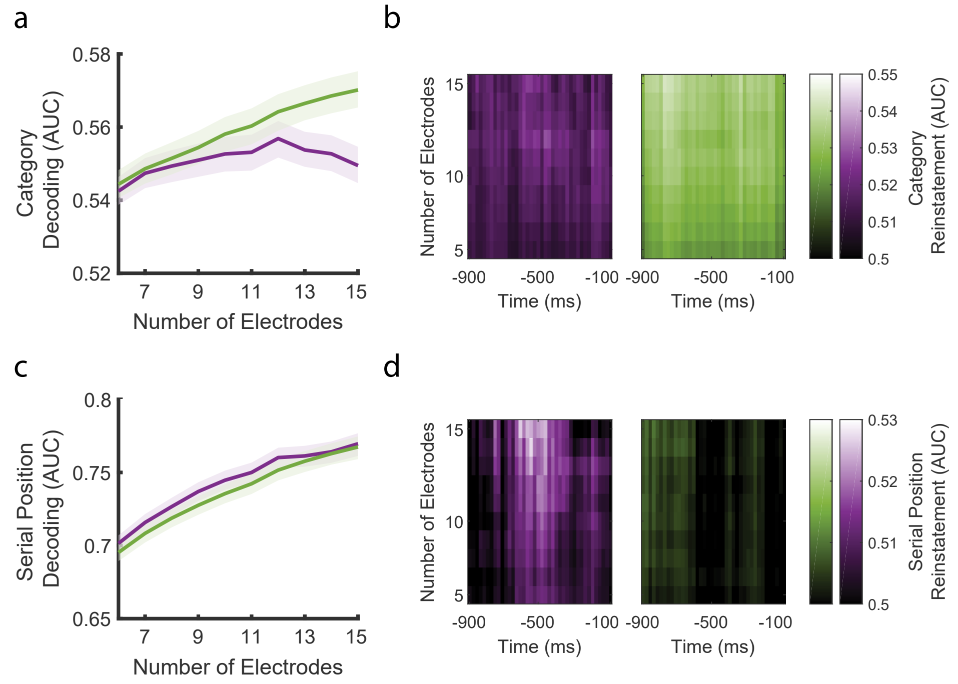


**Supplementary Figure 4  Decoding accuracy increases with electrode coverage. a**, Line plots depict the average category decoding as a function of the number of electrodes sampled from the PM (purple) and AT (green) networks. Shaded regions denote standard error of the mean ($n = 69$ subjects). **b**, Each map depicts category reinstatement in the moments leading up to item recall, within each network. Changes in serial position decoding during encoding (**c**) and associated reinstatement effects (**d**) follow the same plotting convention.
