## Supplementary Figure 5 for "Distinct cortical systems reinstate the content and context of episodic memories"

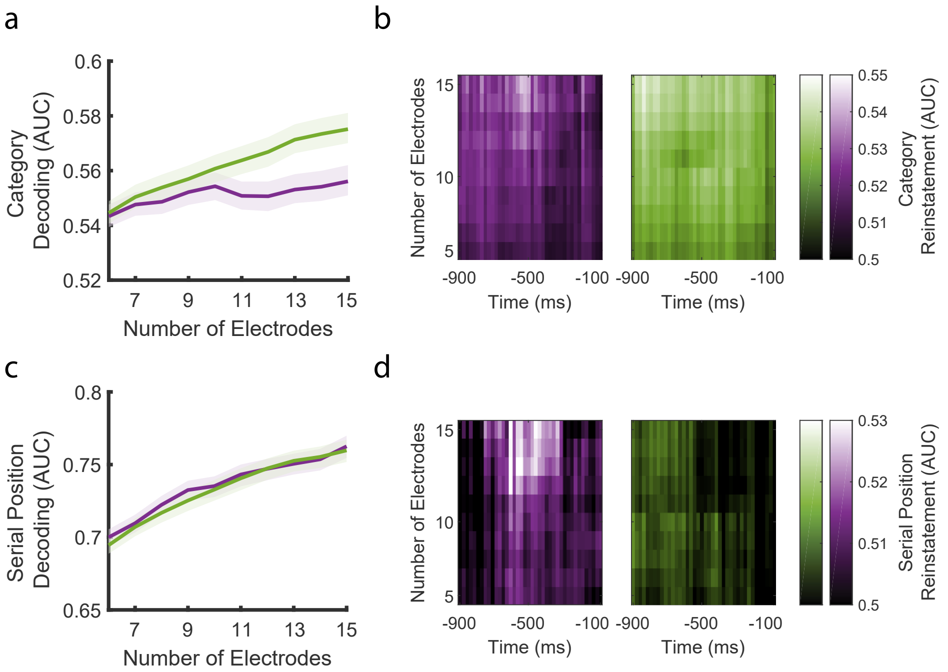


**Supplementary Figure 5 Reinstatement effects are not influenced by epileptogenic brain regions.** Excluding electrodes from the seizure onset zone does not impact category decoding during encoding (**a**) or recall (**b**). The same pattern of results holds for decoding serial position of encoded items (**c**) and reinstatement of related activity (**d**).
