## Supplementary Table 1 for "Distinct cortical systems reinstate the content and context of episodic memories"

**Supplementary Table 1** Patient Demographic Information.

| Subject | Sex | Age (yr.) | Handedness | Seizure Onset Zone (hemisphere) | Epilepsy Duration (yr.) |
|---|---|---|---|---|---|
| 13 | F | 37 | R | Temporal, Occipital (R) | 25 |
| 16 | F | 31 | R | Prefrontal (L) | 30 |
| 21 | M | 39 | R | Temporal, Parietal (B) | 23 |
| 24 | F | 37 | R | Mesial temporal, Parietal, Occipital (R) | 25 |
| 36 | M | 49 | L | Mesial temporal (L) | 34 |
| 42 | F | 28 | L | Frontal, Temporal, Parietal (R) | 22 |
| 45 | M | 51 | R | Mesial temporal (B) | 9 |
| 50 | M | 20 | R | Temporal, Parietal (L) | 6 |
| 60 | F | 37 | R | Prefrontal, Temporal (R) | 22 |
| 65 | F | 34 | R | Temporal (B) | 16 |
| 66 | M | 39 | R | Temporal (R) | 13 |
| 69 | M | 27 | R | Frontal, Parietal (L) | 20 |
| 75 | M | 50 | R | Undetermined | 2 |
| 83 | F | 49 | R | Mesial temporal (L) | 2 |
| 89 | M | 36 | L | Temporal, Prefrontal (R) | 31 |
| 92 | M | 45 | R | Hippocampal (B) | 5 |
| 105 | M | 25 | R | Parietal (R) | 8 |
| 106 | M | 27 | R | Undetermined | 11 |
| 107 | M | 25 | R | Occipital (L) | 24 |
| 108 | F | 24 | R | Hippocampal, Prefrontal (L) | 20 |
| 111 | M | 20 | R | Temporal, Parietal, Occipital (L) | 6 |
| 114 | F | 32 | A | Posterior temporal, Parietal (L) | 10 |
| 127 | F | 40 | R | Temporal, Parietal, Occipital (R) | 11 |

| | | | | | |
|---|---|---|---|---|---|
| 131 | M | 24 | R | Temporal, Parietal (R) | 10 |
| 135 | M | 48 | R | Parietal, Occipital (L) | 39 |
| 138 | M | 42 | R | Mesial temporal, Parietal (B) | 18 |
| 141 | F | 45 | R | Temporal (B) | 38 |
| 157 | M | 22 | R | Amygdala, Frontal, Parietal (R) | 7 |
| 158 | F | 45 | R | Anterior temporal (L) | 40 |
| 163 | M | 46 | R | Mesial temporal (L) | 20 |
| 167 | M | 33 | R | Temporal, Paracentral (L) | 19 |
| 170 | M | 21 | R | Temporal (L) | 12 |
| 174 | M | 29 | R | Temporal, Parietal (L) | 2 |
| 184 | M | 42 | R | Temporal, Parietal (R) | 41 |
| 186 | M | 28 | | Temporal (B) | 15 |
| 187 | F | 52 | R | Parietal, Paracentral | 47 |
| 188 | F | 26 | R | Temporal (R) | 2 |
| 191 | M | 19 | R | Parietal (R) | 7 |
| 192 | M | 28 | R | | 22 |
| 201 | M | 37 | R | Mesial temporal (R) | 3 |
| 202 | F | 30 | R | Prefrontal, Temporal (R) | 18 |
| 212 | M | 47 | R | Frontal, Parietal (R) | 3 |
| 217 | M | 37 | R | Mesial temporal (L) | 35 |
| 227 | M | 33 | R | Parietal (R) | 20 |
| 230 | F | 56 | R | Precentral (L) | 16 |
| 235 | M | 49 | L | Mesial temporal, Prefrontal (L) | 47 |
| 240 | F | 38 | R | Mesial temporal (B) | 3 |
| 260 | F | 57 | R | Mesial temporal, Temporal, Parietal (L) | 49 |
| 264 | F | 53 | R | Mesial temporal (L) | 5 |
| 266 | F | 48 | L | Mesial temporal (L) | 27 |

| | | | | | |
|---|---|---|---|---|---|
| 271 | M | 37 | R | Mesial temporal (R) | 4 |
| 275 | M | 42 | R | Prefrontal, Parietal (R) | 32 |
| 277 | M | 34 | R | Parietal, Paracentral (R) | 14 |
| 278 | F | 22 | R | Parietal (R) | 16 |
| 279 | F | 58 | R | Mesial temporal, Prefrontal (L) | 18 |
| 286 | F | 58 | R | Insular, Parietal (L) | 47 |
| 290 | F | 19 | R | Mesial temporal, Prefrontal, Parietal (L) | 9 |
| 302 | M | 48 | R | Mesial temporal, Temporal, Occipital (L) | 6 |
| 303 | F | 63 | R | Mesial temporal, Temporal, Prefrontal (L) | 21 |
| 313 | M | 22 | R | Parietal, Paracentral (R) | 21 |
| 315 | M | 22 | R | Mesial temporal (L) | 3 |
| 317 | M | 36 | R | Mesial temporal (L) | 15 |
| 320 | M | 57 | R | Mesial temporal (R) | 37 |
| 328 | F | 28 | R | Temporal, Prefrontal (L) | 27 |
| 330 | F | 38 | R | Cingulate (B) | 15 |
| 334 | F | 38 | R | Mesial temporal, Temporal, Parietal (B) | 15 |
| 337 | M | 55 | R | Mesial temporal, Temporal, Parietal, Prefrontal (L) | 18 |
| 354 | M | 28 | R | Mesial temporal, Prefrontal (L) | 14 |
| 361 | F | 31 | | Mesial temporal, Insular, Temporal (B) | 16 |
